# Sablefish (*Anoplopoma fimbra* Pallas, 1814) plasma biochemistry and hematology reference intervals including blood cell morphology

**DOI:** 10.1101/2021.02.01.429130

**Authors:** Carla B. Schubiger, M. Elena Gorman, Jennifer L. Johns, Mary R. Arkoosh, Joseph P. Dietrich

## Abstract

Plasma biochemistry and hematology reference intervals are integral health assessment tools in all medical fields, including aquatic animal health. As sablefish (*Anoplopoma fimbria*) are becoming aquaculturally and economically more important, this manuscript provides essential reference intervals (RI) for their plasma biochemistry and hematology along with reference photomicrographs of blood cells in healthy, fasted sablefish. Blood cell morphology can differ between fish species. In addition, blood cell counts and blood chemistry can vary between fish species, demographics, water conditions, seasons, diets, and culture systems, which precludes the use of RI’s from other fish species. For this study, blood was collected for plasma biochemistry and hematology analysis between June 20 and July 18, 2019, from healthy, yearling sablefish, hatched and reared in captivity on a commercial diet. Overnight fast of 16-18 hours did not sufficiently reduce lipids in the blood, which led to visible lipemia and frequent rupture of blood cells during analysis. Therefore, sablefish should be fasted for 24 to 36 hours before blood is collected to reduce hematology artifacts or possible reagent interference in plasma biochemistry analysis. Lymphocytes were the most dominant leukocytes (98%), while eosinophils were rare, and basophils were not detected in sablefish. Neutrophils were very large cells with Döhle bodies. In mammals and avian species, Döhle bodies are usually signs of toxic change from inflammation, but no such association was found in these fish. In conclusion, lipemia can interfere with sablefish blood analysis, and available removal methods should be evaluated as fasting for up to 36 h might not always be feasible. Also, more studies are required to establish RI for different developmental stages and rearing conditions.

## Introduction

Sablefish (*Anoplopoma fimbria*), colloquially known as butterfish or black cod, are a highly valuable and fast-growing demersal species native to the continental slope of the North Pacific Ocean. Between 2009 and 2019, U.S. landing values of wild sablefish have ranged between $89.1 and $184.6 million (1). Wild sablefish populations have historically declined in the late 1900s due to massive harvest by international fishing fleets, and as a result, the catch is highly regulated to stabilize wild populations (2–5). National and international market demands are projected to continuously increase while harvest of wild sablefish will likely remain limited, resulting in increased interest in an aquaculture industry to meet sablefish demand.

Several research efforts and commercial projects are underway to establish sablefish brood-stock, larval rearing techniques, feeds, as well as the development of net-pen and landlocked recirculating aquaculture systems (RAS) (6–12). Investing in this emerging aquaculture industry brings new opportunities to aquaculture communities and a socioeconomically sensitive approach to satisfy market demands and create jobs beyond what limited catch permits can achieve (13,14).

High stocking density within any culture system increases disease pressure and is a ubiquitous challenge for producers and aquatic veterinarians in charge of maintaining the health status of valuable finfish (15). Research efforts are ongoing to mitigate disease pressure in these culturing systems (16,17). However, little is known about normal physiological health parameters of sablefish. This knowledge gap leaves professionals with insufficient tools to support the health needs of this industry.

Blood parameters are essential indicators of fish health and normal physiology. However, the blood is a complex physiological system, and parameters are sensitive to intrinsic and extrinsic factors, including, in part: fish species, developmental and maturation stage, sex, water chemistry, temperature, nutrition, and management systems (18,19). For example, a previous study evaluated increasing ammonium chloride concentrations on cultured sablefish and identified reduced red blood cell counts, decreased calcium and magnesium levels, and increased liver enzymes (20). Therefore, to support a robust aquaculture industry, studies like the one presented are crucial for establishing reliable reference tools and baselines for each fish species cultured in a commercial system.

This study’s objective was to develop diagnostic references for normal ranges (reference intervals; RI) of plasma biochemistry and hematology values and to characterize blood cell morphology of healthy cultured sablefish. Producers and aquaculture healthcare professionals will be able to use these RI as a guideline and a diagnostic tool.

## Material and methods

Mixed-sex sablefish were obtained from a captive brood-stock and rearing program conducted by the National Oceanic and Atmospheric Administration’s (NOAA) Northwest Fisheries Science Center (NWFSC) at the Manchester Research Station (Port Orchard, WA). The fish were reared for one year in land-based tanks with artificial recirculating seawater at the NWFSC (Seattle, WA) and transferred to NOAA’s Newport Research Station (Newport, OR) on April 18, 2019. At the start of holding, the fish had an average weight of 186 g and grew to an average weight of 440 g on the final day of sampling. In Newport, the sablefish were reared in a 2-m diameter fiberglass tank in ca. 1500 L, with flow-through seawater at ca. 10-14.5 °C. The salinity ranged from 29 to 33.9 ppt and dissolved oxygen ranged from 6.9 to 9.6 mg/L. The indoor tank was lit from above with a combination of natural (skylights) and artificial lighting. The artificial lighting was on a daily timer, providing 9 hours of light, turning on after sunrise, and turning off before sunset. Internal air temperatures ranged from 17.9 to 23.9 °C. Fish were fed BioOregon’s Biobrood (4 mm) diet at least twice per day five days per week, at approximately 1% of body weight. This diet consists of marine fish oils and fishmeal protein. The feed’s composition, per manufacturer, includes 48% minimum protein; 20% minimum oil; 8.5% maximum moisture; 1.0% maximum fiber; 11% maximum ash; 18.2 MJ/kg digestible energy; and 60 ppm astaxanthin (a carotenoid). The sablefish were visually inspected daily Monday through Friday. There were no mortalities or signs of illness observed in the sablefish throughout the duration of the study.

### Preparation of reference animals, sample collection and handling

Pre-analytical procedures were consistent with methodologies applicable to health assessments in aquatic medicine (21). In brief, fish were fasted for 24-36 hours prior to terminal blood collection to avoid lipemia. Two fish were netted and transferred to 19-L buckets containing 500 mg/L tricaine methanesulfonate (MS-222) for euthanasia. After the movement of the operculum stopped, fish were weighed and measured and placed in dorsoventral orientation. Blood was sampled within 3 minutes of euthanasia from the caudal vein. 25 gauge (25 G X 5/8), heparinized needles were inserted at the midline approximately 1-2 cm caudally of the anal fin and advanced craniodorsally at an approximately 60-degree angle to just ventral of the spine. Approximately two to four mLs of blood per fish were collected in the heparinized syringes. Immediately after collection, blood smears were prepared, and Fisherbrand™ non-heparinized microhematocrit capillary tubes (Fisher Scientific, Carlsbad, CA) were filled to determine hematocrit and total plasma protein.

An aliquot of heparinized blood (1.5 mL) was centrifuged at 1200 x g at 14 °C for 10 minutes. Plasma was harvested and transferred to a sterile 2 mL empty tube. Heparinized whole blood and plasma samples from each fish were stored in a chilled container and transported within 2-4 hours to Oregon State University Veterinary Diagnostic Laboratory in Corvallis, Oregon.

Blood was collected from a total of 87 sablefish. However, data from 6 fish (sampled on June 20 and July 2, 2019) were excluded from analysis due to grossly high lipid content in the blood that could potentially cause aberrant biochemical and hematological analyses and skewing statistical results. Therefore, reference intervals were ultimately determined from a total of 81 sablefish of the same lineage, age, and environmental conditions. Fish were collected on June 24 (2 fish), 25 (3 fish), 26 (3 fish), and July 5 (1 fish), 10 (32 fish), and 17 (40 fish), 2019. This study was permitted by the animal care and use committee at Oregon State University (ACUP 5154).

### Hematology and Biochemistry Methods

Heparinized whole blood and plasma samples were immediately processed upon receipt by the Veterinary Diagnostic Laboratory at the Oregon State University Carlson College of Veterinary Medicine in Corvallis, Oregon. The lab evaluated the biochemical and hematologic parameters.

Plasma biochemical profiles were selected based upon prior studies in other fish species (22–24). Biochemical analysis was performed via the Beckman Coulter AU480 automated chemistry analyzer (Beckman Coulter Inc., Brea, CA, USA) and reported in conventional units. The conventional units were then converted to the International System of Units (SI units) (25). The biochemical parameters analyzed are listed in Table 1.

**Table 1.** Plasma biochemistry parameters.

| Plasma Biochemistry Parameter & Unit of Measurement | Description |
| --- | --- |
| Albumin (Alb) | Protein synthesized by the liver, important for oncotic pressure in blood and carrier function of hormones, enzymes, vitamins, etc. |
| Globulins (Glob) | Proteins synthesized in liver and immune system, important in immune response, liver function, blood clotting, etc. |
| Total Protein (TP) | Albumin and globulins combined |
| Glucose | Simple sugar important energy source |
| Cholesterol | Lipid synthesized in cells, serves as component of cell membranes and precursor of steroid hormones and vitamin D. |
| Creatinine | Breakdown product from muscle metabolism, excreted by kidneys |
| Urea Nitrogen (BUN) | Waste product produced in liver from protein digestion, excreted by kidneys |
| Total Bilirubin | Breakdown product of red blood cells, excreted via liver, kidneys, and gastrointestinal tract |
| Alanine Aminotransferase (ALT) | Formerly called serum glutamate-pyruvate transaminase (SGPT), enzyme present in normal liver and muscle cells |
| Aspartate Aminotransferase (AST) | Transport enzyme in amino acid metabolism, biomarker for liver health, also known as serum glutamic oxaloacetic transaminase (SGOT) |
| Creatine Kinase (CK) | Enzyme marker for muscle damage |
| Sodium (Na) | Electrolyte to maintain fluid levels and pH |
| Potassium (K) | Electrolyte involved in muscle and nerve activity, fluid levels and more |
| Chloride (Cl) | Electrolyte involved in blood volume and pressure, and pH |
| Calcium (Ca) | Mineral in bone; levels can alter with bone, kidney, endocrine, nutritional or toxic disorders |
| Phosphorus (P) | Mineral in bones and teeth, involved in nerve signaling and muscle contractions, abnormal levels indicative of kidney, liver, or bone diseases |
| Total CO<sub>2</sub> | Indicator of bicarbonate in serum, involved in pH regulation |
| Anion Gap | Indicator of the amount of “unmeasured” ions (usually anions) in blood, increases are seen with lactic acidosis, ketosis and renal disease, decreases are usually due to loss of albumin |

Measurement of all hematologic parameters was performed manually within 24 hours of receipt. Heparinized whole blood was utilized to assess packed cell volume (PCV), total protein (TP), and manual cell counting. PCV was obtained by the collection of whole blood into hematocrit tubes and measurement post-5-minute centrifugation. Total plasma protein was evaluated by a refractometer (Reichert, Depew, NY, USA). Manual cell counts were performed by two board-certified clinical pathologists (EG and JJ). Leukocyte and thrombocytes were stained by transferring 20 µL whole blood into 1.98 mL Natt-Herrick Solution (Vetlab supply, Palmetto Bay, FL, USA) for a 1:100 dilution. The mixture was allowed to incubate for 20-30 minutes at room temperature then counted via Neubauer hemocytometer (Bright-Line, Hausser Scientific, Horsham, PA) using a standard protocol (22,26,27). Leukocytes were counted in nine primary squares, and the total was divided by nine and multiplied by 100 (dilution factor). Thrombocyte counts were obtained by counting thrombocytes in the center main-square of the hemocytometer then multiplying by 1000.

Blood smears were submitted on glass slides and stained with Modified Wright’s stain (Thermo Fisher Scientific) via an automated method (HemaTek 2000, Siemens Healthcare, Erlangen, Germany). Leukocytes were identified based upon characteristics described for other related marine species (28,29), and morphologic identity was initially agreed upon by both pathologists (EG and JJ). Leukocyte differential counts were performed on 300 cells in each slide and converted to absolute WBC counts. The differential counts were performed by an individual pathologist (EG) to minimize inter-observer variability.

### Reference interval calculations

Blood values were collected into Excel spreadsheets and reference intervals calculated with the macro add-in Reference Value Advisor V 2.1 software (30). The software follows recommendations and guidelines by the Clinical and Laboratory Standards Institute (CLSI), the National Committee for Clinical Laboratory Standards (NCCLS), and the International Federation of Clinical Chemistry (IFCC). The software determined that the sample size was large enough to compute nonparametric reference intervals and also determined the 90% confidence intervals of the limits of the nonparametric reference intervals by a bootstrap method (30,31).

## Results

### Plasma biochemistry reference intervals

The reference intervals for measured plasma biochemistry parameters in the 81 sufficiently fasted sablefish is presented in Table 2. Overnight fasting of 16-18 hours was not sufficient to reduce visible lipemia of the blood in three of the sablefish sampled on June 20, 2019 (animals #32, #33, #34). Also, an unscheduled blood collection was performed on July 2, 2019, with fish that were fasted for only 16-18 hours leading again to grossly lipemic samples (animals #47, #48, #49). Consequently, the protocol was adjusted to increase the fast time to 24-36 hours. Therefore, six of the 87 animals were excluded from the reference interval calculations due to the severely lipemic condition of the plasma samples. This condition displayed as visible turbidity in some samples (S1 Fig) and severe blood cell rupture in others.

**Table 2.** Reference intervals for plasma biochemistry parameters of fasted sablefish.

that were fasted 36 to 48 hours. Data were collected in conventional units and converted into SI units according to (25).
| Parameter; n = 81 | SI units | Conventional units |
| --- | --- | --- |
| Albumin (Alb) | 6.1 – 13 g/L | 0.61 – 1.3 g/dL |
| Globulin (Glob) | 19.1 – 31.9 g/L | 1.91 – 3.19 g/dL |
| Total Protein (TP) | 25.2 – 44.9 g/L | 2.52 – 4.49 g/dL |
| Glucose | 0.95 – 4.91 mmol/L | 17.1 – 88.4 mg/dL |
| Cholesterol | 6.34 – 12.48 mmol/L | 245.2 – 482.5 mg/dL |
| Creatinine | 0 – 140.46 µmol/L | 0 – 1.59 mg/dL |
| Urea Nitrogen (BUN) | 0.07 – 0.14 mmol/L | 2.0 – 4.0 mg/dL |
| Total Bilirubin | 1.71 – 3.42 µmol/L | 0.1 – 0.20 mg/dL |
| Alanine Aminotransferase (ALT, SGPT) | 0 – 17.8 U/L | 0 – 17.8 U/L |
| Aspartate Aminotransferase (AST, SGOT) | 0 – 1.88 µKat/L | 0 – 110.4 U/L |
| Creatine Kinase (CK) | 0.09 – 1.49 µKat/L | 5.1 – 87.4 U/L |
| Sodium (Na) | 168.0 – 179.9 mmol/L | 168.0 – 179.9 mEq/L |
| Potassium (K) | 1.2 – 3.86 mmol/L | 1.2 – 3.86 mEq/L |
| Chloride (Cl) | 142.0 – 153.0 mmol/L | 142.0 – 153.0 mEq/L |
| Calcium (Ca) | 2.63 – 3.35 mmol/L | 10.51 – 13.38 mg/dL |
| Phosphorus (P) | 4.17 – 8.41 mmol/L | 12.92 – 26.06 mg/dL |
| Total CO<sub>2</sub> | 4.64 – 8.70 mmol/L | 4.64 – 8.70 mEq/L |
| Anion Gap | 18.1 – 27.0 mmol/L | 18.1 – 27.0 mEq/L |

### Hematology Reference Intervals

Hematologic parameters, including packed cell volume (PCV), plasma protein (PP), and cell counts, were collected manually from the 72 fish sampled on July 10 and 17, 2019. Thrombocyte counts were only available from 60 animals due to early technical difficulties with staining. All data are presented in Table 3.

**Table 3.** Cell blood counts (CBC) were available from 72 animals, with the exception that thrombocyte counts were only available from 60 animals.

legend: PCV, PP, and blood cell counts were manually performed from fish sampled on July 10 and 17, 2019.
| Parameter | n = 72 (*n = 60) |
| --- | --- |
| Packed Cell Volume (PCV) | 0.36 – 0.62 L/L |
| Plasma Protein (PP) | 43.0 – 79.2 g/L |
| Thrombocytes * | 92.1 – 357.48 x10<sup>9</sup> /L |
| White Blood Cell (WBC) | 4.33 – 33.08 x10<sup>9</sup> /L |
| % Neutrophils | 1.8 – 23.4 % |
| Neutrophils | 0.08 – 7.32 x 10<sup>9</sup> /L |
| % Lymphocytes | 66 – 96 % |
| Lymphocytes | 3.76 – 24.25 x 10<sup>9</sup> /L |
| % Monocytes | 0 – 12.1 % |
| Monocytes | 0 – 2.92 x 10<sup>9</sup> /L |
| % Eosinophils | 0 - 2.2 % |
| Eosinophils | 0 – 0.49 x 10<sup>9</sup> /L |
| % Basophils | 0 % |
| Basophils | 0 x 10<sup>9</sup> /L |

Morphology of blood cells was similar to those noted in other fish species (28,29,32). Erythrocytes are oval with a round to ovoid nuclei (Fig 1a,b,c). Polychromatic erythrocytes were commonly found in sablefish, which indicates an active turnover of erythrocytes (Fig 1a). Additionally, fish have a thin barrier between hematopoietic tissue and the vasculature; hence, immature erythroid cells were common and may comprise more than 10% of all erythrocytes in circulation (28,32) (Fig 1c). Immature erythrocytes are smaller and more round than mature erythrocytes with basophilic cytoplasm (Fig 1a). Mitotic figures were not readily identified in blood smears of this population.

**Fig 1a.**
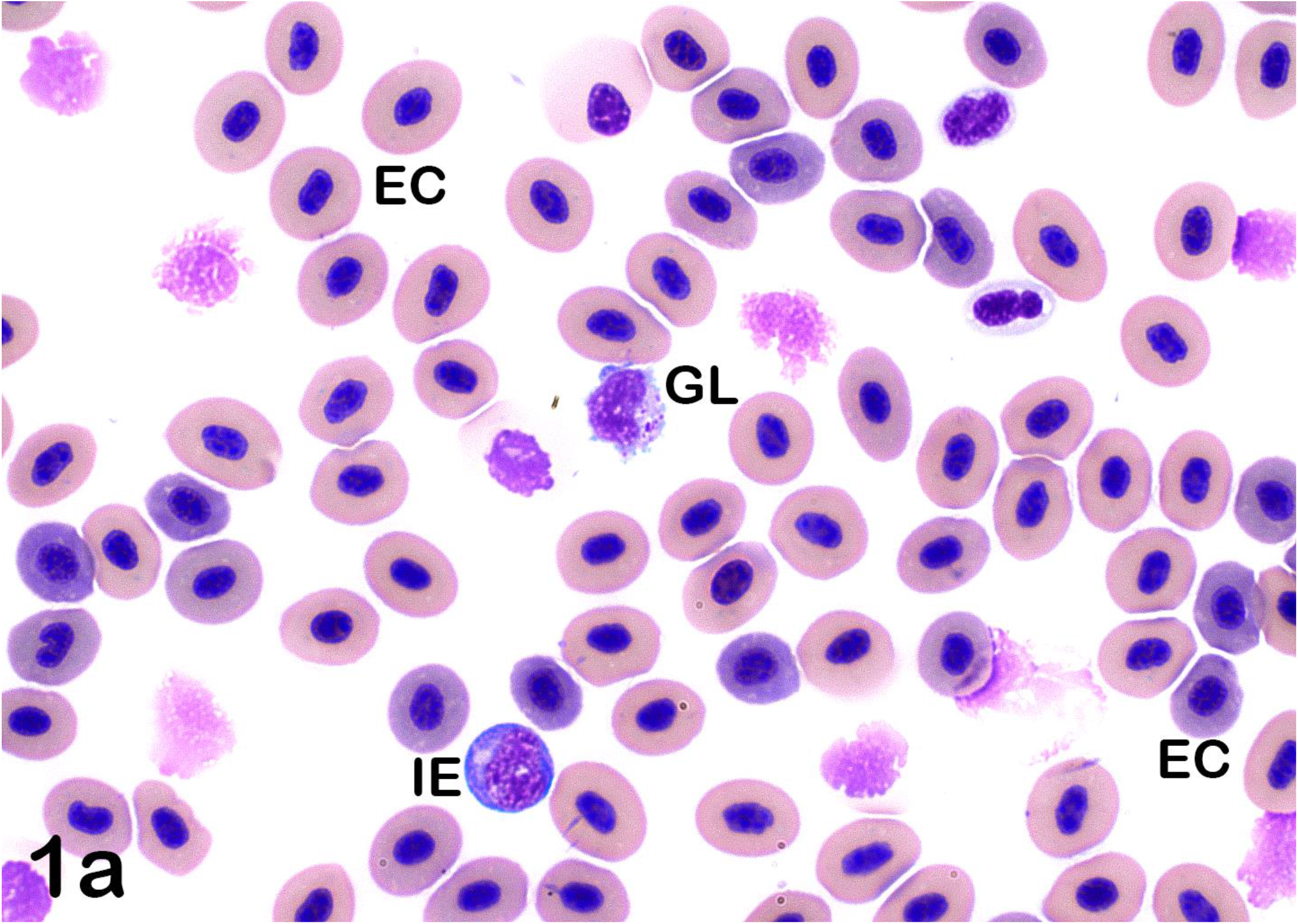
Lymphocytes: Granular lymphocytes. Granular lymphocyte (GL), surrounded by rounder, basophilic immature erythrocytes (IE) that are smaller than the more mature erythrocytes (EC). Overall erythrocytes are in various stages of maturity contributing to a distinct polychromatic appearance, ranging from slightly eosinophilic to basophilic (Wright’s stain x100 objective).

**Fig 1b.**
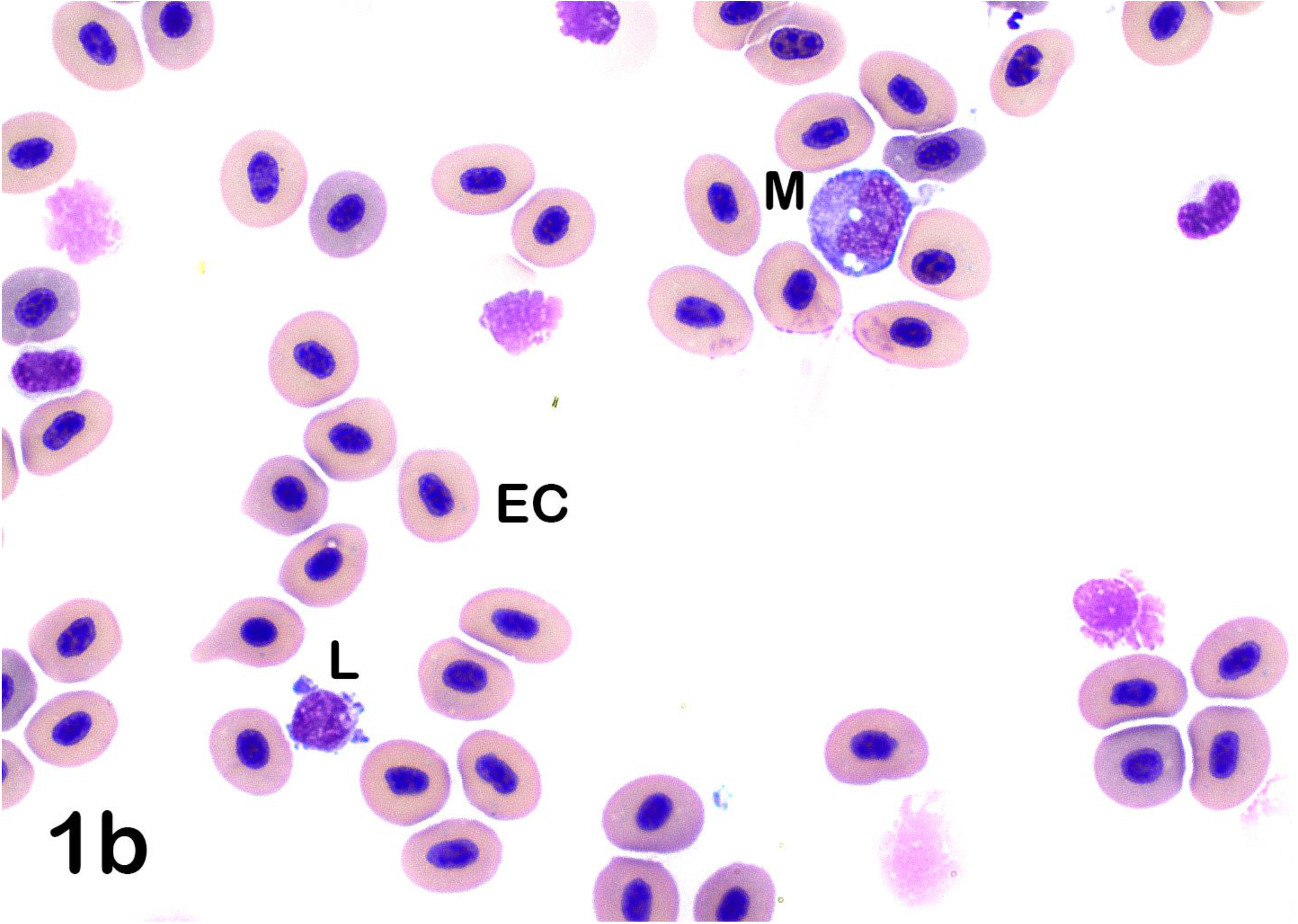
Lymphocytes: Mature lymphocyte and monocyte. Mature lymphocyte (L) and monocyte (M) with typical large oval eccentrically-positioned nucleus and a moderate amount of basophilic cytoplasm that contains three discrete cytoplasmic vacuoles. Mature lymphocytes are characterized as round cells with large, round nucleus with condensed chromatin and scant basophilic cytoplasm. Lymphocytes often display blebbing of the cell borders (Wright’s stain x100 objective).

**Fig 1c.**
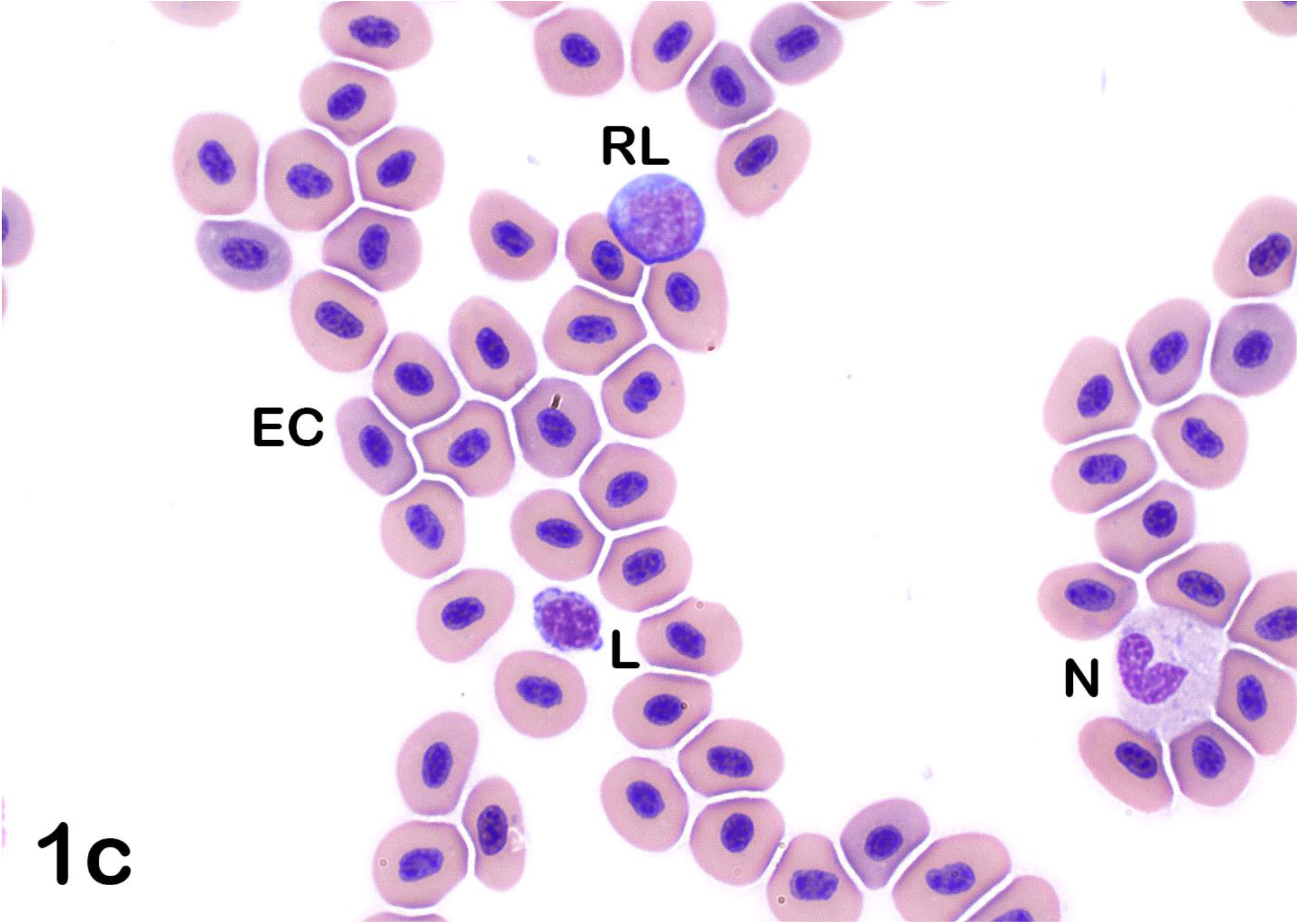
Lymphocytes: Reactive lymphocyte. Reactive lymphocyte (with Golgi body left to the nucleus, RL), small lymphocyte (L) with blebbing and neutrophil (N). Reactive lymphocytes are large cells, with a more open chromatin pattern and increased amounts of basophilic cytoplasm (Wright’s stain x100 objective).

In sablefish, lymphocytes predominated and comprised up to 98% of the total leukocytes present in healthy individuals. Lymphocyte morphology was similar to that of other species, and lymphocytes were identified as round cells with high nucleus to cytoplasm ratios (N:C ratios), round nuclei and condensed chromatin (Fig 1a-c). Cell borders of lymphocytes were frequently blebbed (bulging plasma membrane) (Fig 1c). Reactive lymphocytes were also similar to those in other species, with cells being larger than mature lymphocytes, more open chromatin patterns, and increased basophilic cytoplasm. Occasional Golgi bodies were noted (Fig 1c). Additionally, granular lymphocytes were occasionally found (Fig 1a). Neutrophils were one of the largest leukocytes identified in sablefish (Fig 1c, 2, 3). Neutrophils were oval with oval, clefted, or segmented nuclei surrounded by moderate to abundant amounts of slightly eosinophilic cytoplasm. Most neutrophils contained pale blue cytoplasmic inclusions consistent with Döhle bodies. These bodies are often signs of inflammation or bacterial infection in mammals and birds, but appeared to have no association with other toxic changes in these fish (Fig 1c, 2, 3) (33–35).

**Fig 2.**
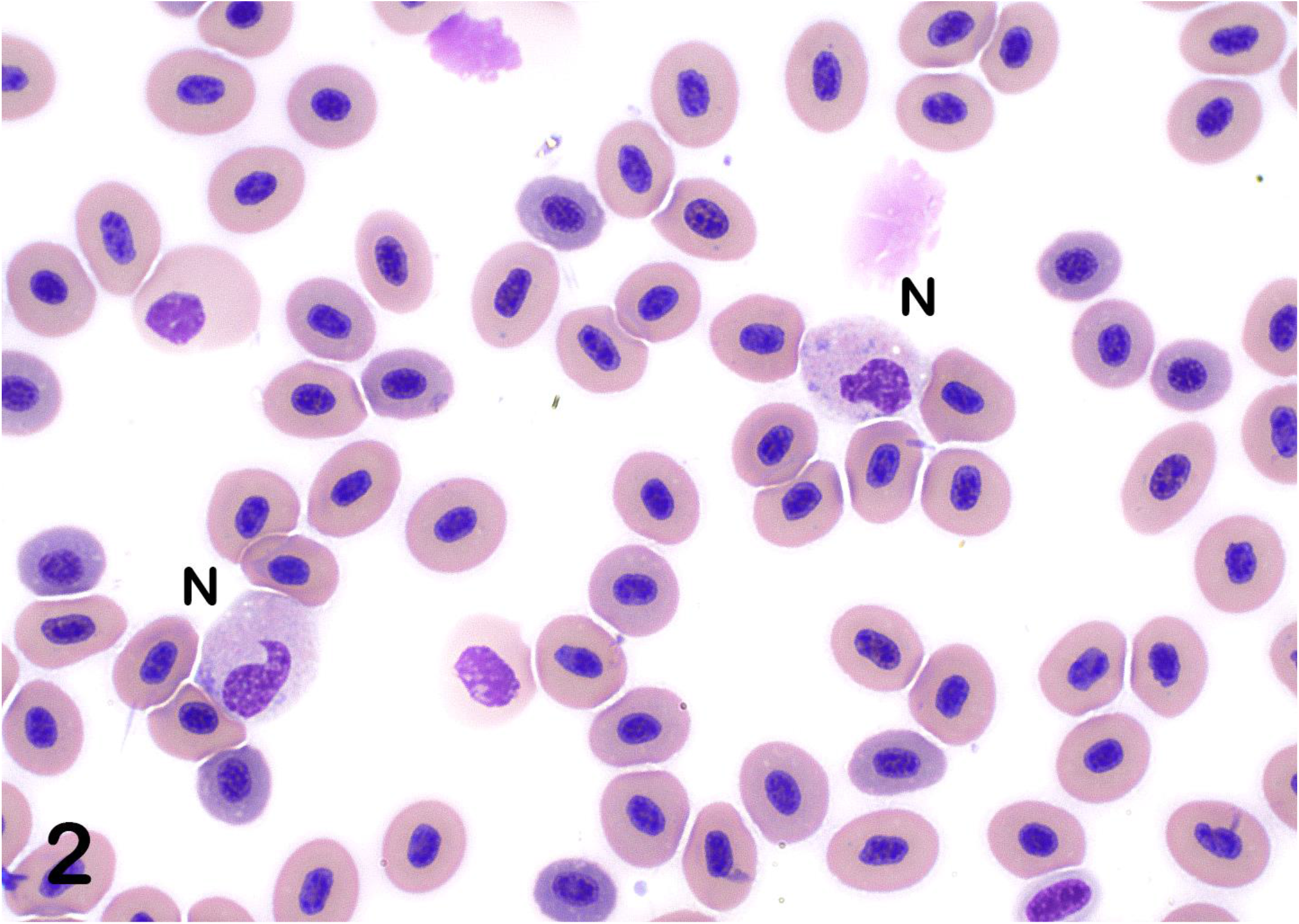
Neutrophils. Neutrophils (N) are large leukocytes with large oval, clefted, or segmented nuclei and moderate to large amounts of slightly eosinophilic cytoplasm that often contains pale blue cytoplasmic inclusions consistent with Döhle bodies (Wright’s stain x100 objective).

**Fig 3.**
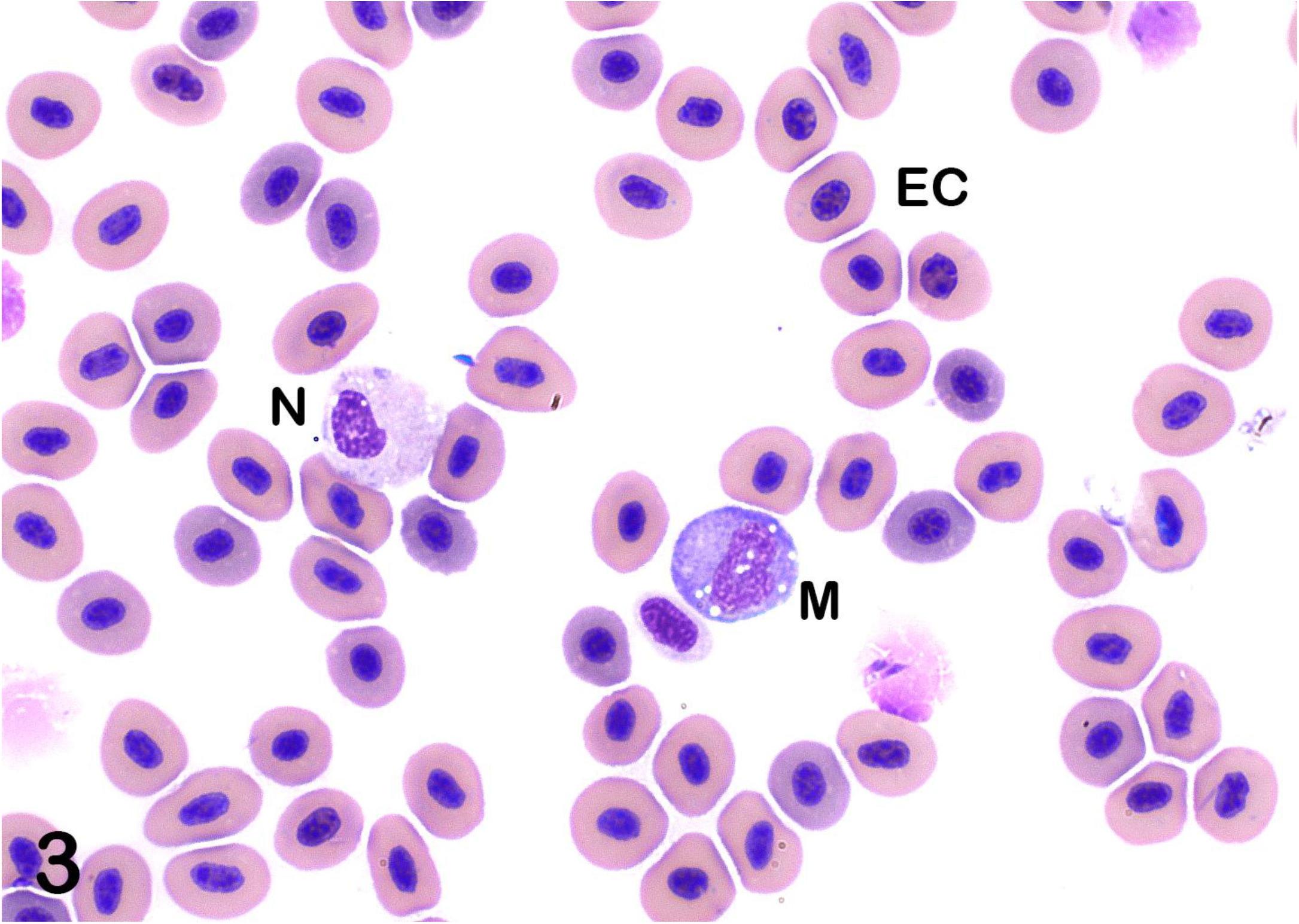
Neutrophil (N) and Monocyte (M). Note similar size between these cells. Mature erythrocytes (EC) also identified (Wright’s stain x100 objective).

Monocytes were larger than lymphocytes and similar in appearance to monocytes in other non-mammalian species (Fig 1b, 3, 4). Monocytes contained oval eccentrically-positioned nuclei; amoeboid nuclear shapes were less frequent. Monocytes had small to moderate amounts of variably basophilic cytoplasm and frequently contained discrete cytoplasmic vacuoles (Fig 1b, 3, 4). Cell borders were distinct and non-blebbed, making them easily distinguishable from lymphocytes. Eosinophils were rare in healthy sablefish. These were small cells with a high N:C ratio (Fig 5). Nuclei of eosinophils were oval and typically eccentric, and cytoplasm contained small, bright eosinophilic granules (Fig 5). Basophils were not found. Thrombocytes were found individually and in small aggregates (Fig 4, 6). Thrombocytes were small and typically oval to elongate in shape. Few round forms were also noted. Nuclei of thrombocytes were pleomorphic, ranging from round to oval to reniform (Fig 4, 6). The cytoplasm was clear with small cytoplasmic granules occasionally seen.

**Fig 4.**
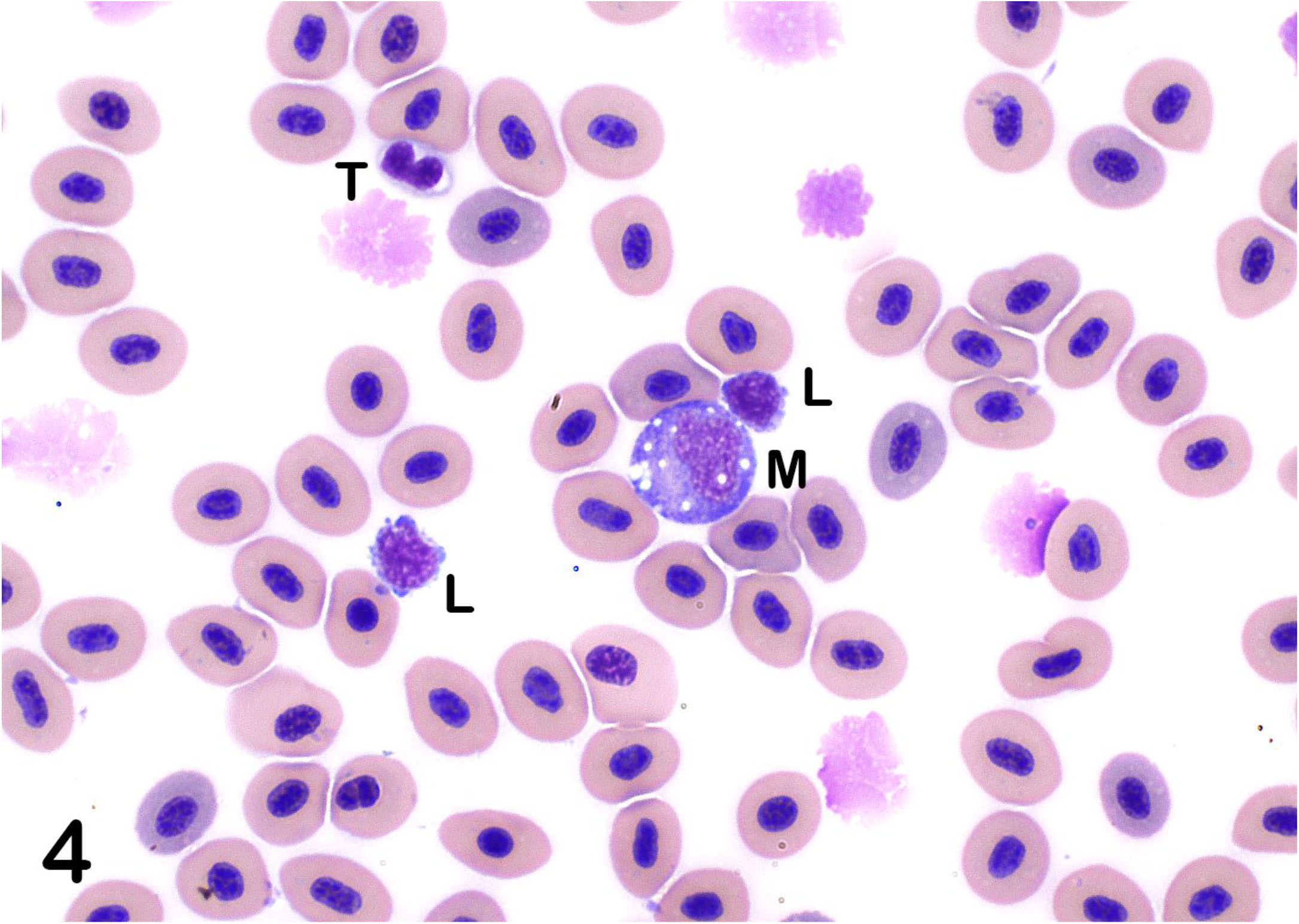
Monocytes. Monocyte (M) contains several, variably sized, discrete cytoplasmic vacuoles and distinct, smooth cell borders, in contrast to the two lymphocytes (L) that display distinct blebbing. Also present is a thrombocyte (T) with a bi-lobbed nucleus (Wright’s stain x100 objective).

**Fig 5.**
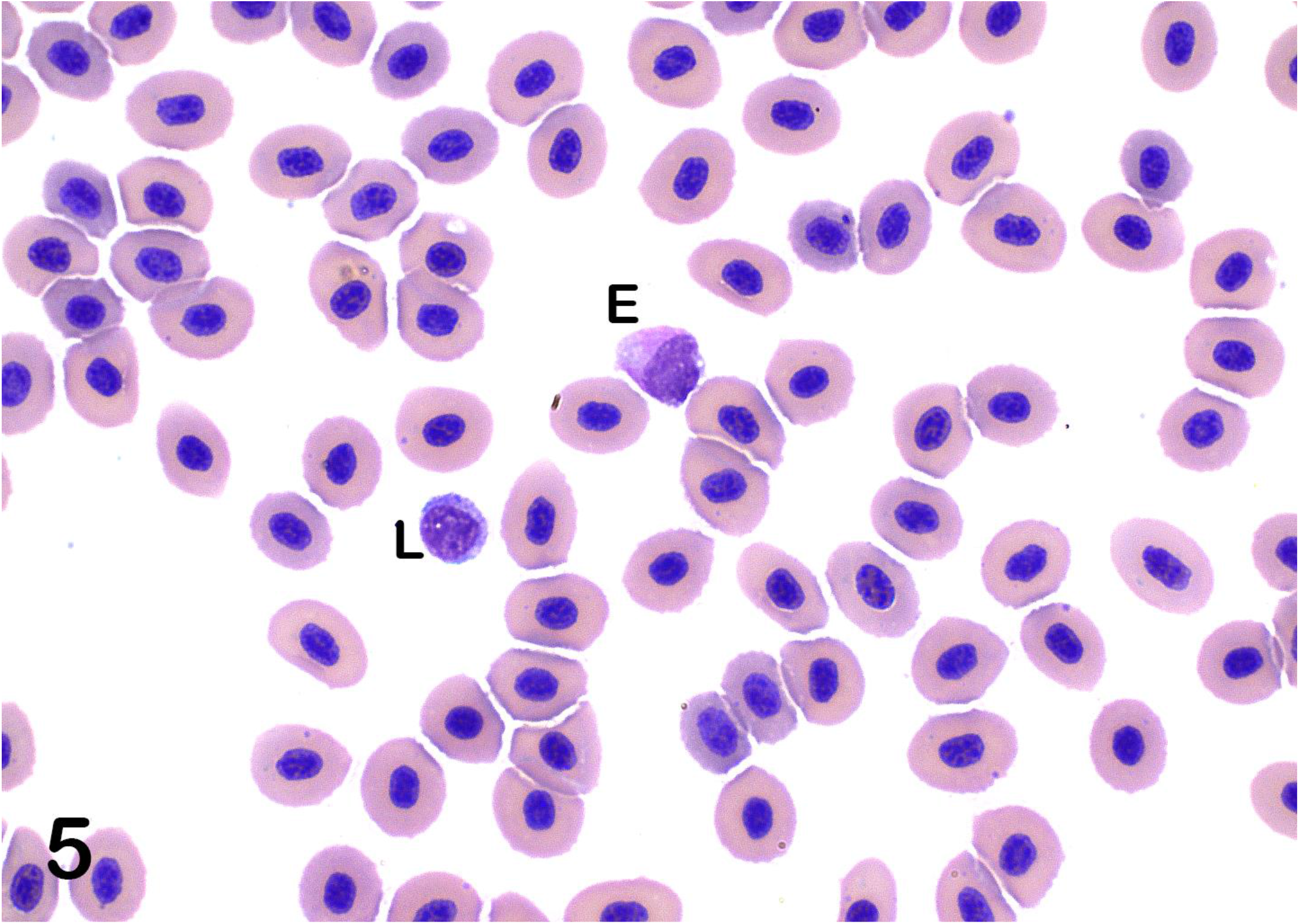
Eosinophils. Eosinophils (E) are slightly larger than mature lymphocytes (L) and have large, ovoid, eccentric nuclei and small amounts of cytoplasm that contains small, bright eosinophilic granules (Wright’s stain x100 objective).

**Fig 6.**
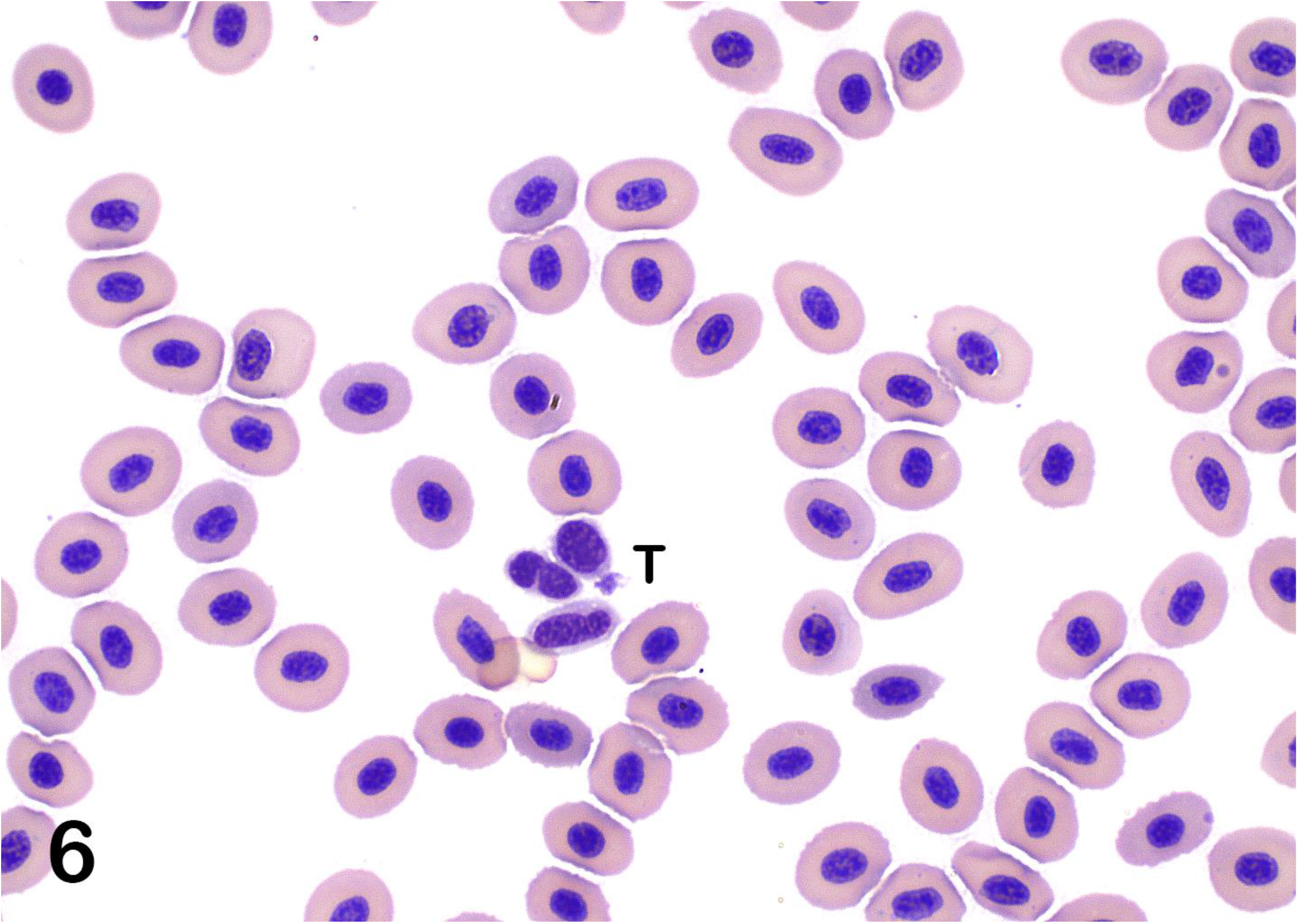
Thrombocytes (T). Thrombocytes are found individually and in small aggregates in well-prepared blood smears. Thrombocytes are small and typically oval to elongate and commonly displaying nuclear pleomorphism with round to oval to reniform nuclei. The cytoplasm of thrombocytes is usually clear with occasional, small cytoplasmic granules (Wright’s stain x100 objective).

## Discussion

This study addresses an urgent need associated with the development of a sustainable and highly profitable emerging aquaculture species. Establishing a sablefish aquaculture industry would contribute to the livelihoods of coastal (and inland) communities along the northern Pacific Ocean. Species-specific biochemistry and hematologic reference intervals (RI) derived from a healthy reference population are crucial for adequate health monitoring of any species, particularly species new to culturing. Management and feeding practices, and disease treatment regimens can be appropriately adjusted in correlation to physiological parameters of the fish determined from health monitoring. Blood cell counts are valuable tools in assessing the general health status of the fish and can alert a health care provider to infections and environmental or management issues, e.g., problems with water quality or nutrition.

An important observation was that fasting sablefish for 24-36 hours was necessary to obtain blood samples with lipid content low enough to not interfere with plasma biochemistry or hematology analysis. High amounts of triglycerides in human blood has been found to significantly interfere with measurements of phosphorus, calcium, creatinine, and total protein due to an interaction with the analyzer and reagents (36,37). Plasma protein measured via a refractometer was predictably increased relative to plasma total protein measured by an automated method as lipids refract light and can falsely increase refractometric measurement of protein. High lipid content is also suspected of causing premature lysis of blood cells (38). If fasting is not possible, lipid removal by centrifugation or lipid extraction with polar solvents is possible (39,40). The Clinical and Laboratory Standards Institute (CLSI) recommends ultracentrifugation, but this procedure is not always a practical option (41). Alternatives that might be more practical for onsite blood diagnostics are dilution of the serum sample or usage of serum blanks (40). However, if blood samples are necessary for diagnostics in disease events, versus routine health checks, lipemic blood samples might be less common as inappetence is often the first clinical sign of problems in a culturing system. For scheduled health checks, extended fasting of at least 24 hours is recommended for sablefish based on the results presented in this study. For our analysis, we excluded samples with grossly visible lipemia (milky, turbid appearance), because of possible interference with biochemistry values (animals #32-34, sampled on June 20) and blood samples with moderate cell rupture that was likely caused by lipemia (animals #47-49, sampled on July 2). To present how these exclusion criteria influenced the plasma biochemistry intervals, the complete data set of all 87 fish is presented as supplemental material (S1 Appendix). The exclusion of these insufficiently fasted fish (16-18 hours) notably widened the glucose range, but decreased the upper RI limits of cholesterol, creatinine, bilirubin, and the anion gap. Changes to other parameters were smaller and often in the decimals. A comparison of all values is presented as supplemental material (S1 Appendix).

The fish used in this study received a commercial diet at 1% of their body weight distributed over at least two feeding events per day. Recent studies recommend feeding sablefish to satiation once every other day to allow for adequate feed conversion and consistent intake of the feed (42,43). Reduced feed intake has been observed after four days of daily feeding, attributed to a prolonged gastrointestinal passage in this species (42) (and personal communication with Ron Johnson). The delayed clearance of blood lipids experienced in this study is likely a reflection of that observation.

The possibility exists that the observed affinity for high blood lipid concentration reflects the general tender consistency of the meat in this species, which gave rise to the colloquial name “butterfish”. However, the energy density of sablefish is relatively low compared with other pelagic fish from the North Pacific, and only low to moderate amounts of the wet mass are attributed to lipids (44). A muscle lipid content of up to 14% has been reported in sablefish with higher amounts of lipids stored in the bones (44–48). Nevertheless, sablefish seem to have a well-adapted fat metabolism. Earlier studies have shown that prolonged starvation of five months in a laboratory setting did not result in significant protein catabolism. Instead, fish survived by utilizing their fat storage over muscle tissue, making sablefish generally well adapted to infrequent meals in the wild (46,49). Sablefish are identified as opportunistic feeders, whose food sources can alter by location and age (50). Older and spawning fish commonly reside in deeper water, while the larvae hatch in the deep and then migrate up, and younger fish are more commonly found in shallower waters (11,50,51). Besides the ability to travel very long distances (an average of 191 km per year (52)), sablefish can also undergo an impressive diel vertical migration in pursuit of some of their vertically migrating prey by ascending to the surface at night and descending to waters of 750 m or greater before sunlight (50). This exposes the animal to stark differences in temperature and pressure. More detailed reasons for these migrations and the intricacy of sablefish lipid physiology are still under further investigation.

Animal husbandry and management can have a substantial impact on blood reference ranges. In a study of blood RI in sunshine bass, the only biochemical reference intervals that were similar in three different holding systems were glucose and cortisol (53). Similarly, previous studies in sablefish describe that tank design and water temperature can influence survival, anatomical deformities, meat quality, and ovarian development (6–8). Moreover, studies found that blood values in sturgeon can differ among hatcheries (24). Stress is reported to alter the fish’s ability for osmoregulation, which will affect electrolyte concentrations in the blood, often resulting in chloride loss and high glucose values (53,54). Fish sampled in this study had a wide range of glucose levels (0.95–4.91 mmol/L, 17.1-88.4 mg/dL). This was mostly driven by noticeably higher glucose levels in the last ten of 32 fish and 40 fish sampled on July 10 and July 17, respectively (S1 Appendix). Most of the glucose measurements were in the upper teens to mid-forties mg/dl (approximately 1-2.5 mmol/L). The later upward trend of glucose levels may be due to the repeated netting of fish from the holding tank during the two to three hours of sampling. Chloride (142-153 mmol/L, 142-153 mEq/L) was below what has been suggested as normal for marine teleosts (165-175 mmol/L, 165-175 mEq/L), but slightly higher than reported in Atlantic salmon (125-143 mmol/L, 125-143 mEq/L) which were regarded relatively stress-resistant, and much higher than reported in Atlantic sturgeon (116-131 mmol/L, 116-131 mEq/L) (24,55,56). These sources would also suggest that the sablefish RI’s of monovalent sodium (168-179.9 mmol/L, 168-179.9 mEq/L) and potassium (1.2-3.86 mmol/L, 1.2-3.86 mEq/L), and bivalent calcium (2.63–3.35 mmol/L, 10.51-13.38 mg/dL) are lower than in other marine teleosts (175-200 mmol/L, 5-10 mmol/L and 3-5 mmol/L, respectively) (24,55). The divergence of biochemical reference intervals in older literature and more recent comprehensive reviews on intrinsic and extrinsic influences on blood chemistry and hematology suggest that further studies that consider fish demographics, feeds, and production systems are warranted (18,19).

Sex can also affect blood reference intervals (55). All-female stocks are preferred in the production of many food fish species, including sablefish because female fish grow larger than males (7,57). All-female stocks can be achieved by mating females with neomales, which are genetic females who express phenotypically male. This can either be achieved by the oral administration of testosterone or by temperature management during the sexually undifferentiated phase early in development (7). The mixed-sex population sampled in the present work warrants further comparison and investigations into the physiology of this species. In addition, the present sablefish blood RI should be compared to blood values of sablefish that are cultured in other production systems, including recirculating aquaculture systems (RAS) and marine net pens, in different age groups, and from progeny that comes from cultured rather than wild, captive brood-stock.

## Acknowledgments

The authors thank Frank Sommers, Ron Johnson, and Rick Goetz of NOAA Fisheries, NWFSC for supplying the cultured sablefish, and are grateful for Annabel Mendoza, Serina Moheed, Brak Moczygemba, and Mary Beth Rew Hicks for animal husbandry and fish necropsy assistance. This study was funded by an Oregon Sea Grant Program Development Award #NA18OAR4170072 awarded to CBS and partially supported by an Oregon State University Agricultural Research Foundation Grant #9026A awarded to CBS.

## Supporting information

**S1 Fig. Lipemic blood**. Photograph of lipemic blood sample from fish #34 on June 20th, 2019, after centrifugation. A packed cell fraction (bottom of tube) is overlaid by a turbid, pale-beige plasma. The turbidity indicates a significant amount of blood lipids.

**S1 Appendix. Raw data: Plasma biochemistry reference intervals**. Data set includes lipemic samples, RI calculation reports for each biochemistry parameter, and a comparison of the RI of 87 fish (with lipemic samples) versus 81 fish (without lipemic samples), plus all conversions from the conventional units to the SI units.

