## Supplemental data Figure S1 for "Sablefish (*Anoplopoma fimbra* Pallas, 1814) plasma biochemistry and hematology reference intervals including blood cell morphology"

**S1 Figure.**

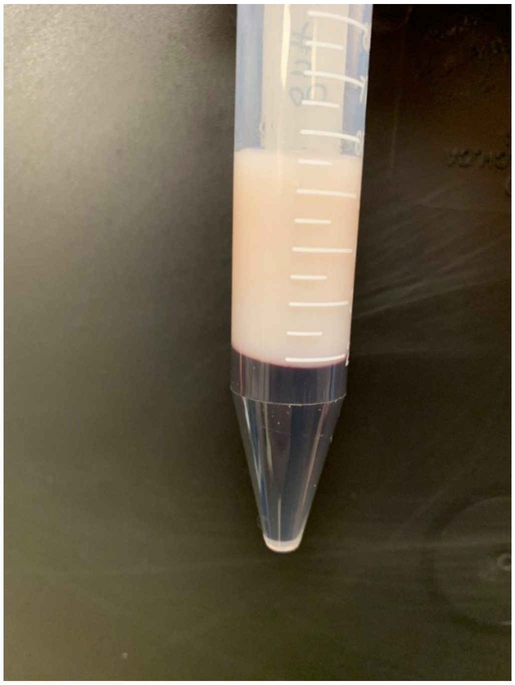
